# Stochastic gene expression drives state switching and phenotypic heterogeneity in RNA repair

**DOI:** 10.64898/2026.11.03.763983

**Authors:** Hollie J. Hindley, Rodrigo Lara Molina, Andrea Y. Weiße

**Affiliations:** Centre for Engineering Biology, School of Biological Sciences, University of Edinburgh, UK; School of Informatics, University of Edinburgh, UK

## Abstract

Bacterial populations often exhibit variability in cellular responses to antibiotics which can affect treatment success and survival outcomes. This heterogeneity suggests that genetically identical cells can occupy different states of susceptibility within the same population. Better understanding of these dynamic responses is essential for determining how cells respond to stress and survive antibiotic-induced damage. In this work, we investigate heterogeneity in antibiotic responses driven by the Rtc RNA repair system in *E. coli*. We develop a stochastic model of Rtc-mediated RNA repair to identify the drivers of heterogeneity at the single-cell level. We observe bimodality in cellular translational capacity and a transient memory in predicted responses to RNA damage, demonstrating how intrinsic stochasticity in Rtc-regulation can lead to phenotypic heterogeneity. Classifying single-cell trajectories, we predict drivers of state switching between the resistant and susceptible states, finding that stochastic expression of the *rtc* regulator determines entry into a resistant cell state, while dilution drives the system back into a susceptible state. These results indicate that heterogeneity in RNA repair emerges from stochastic switching between states, offering a route to understanding how individual cells survive in the presence of translation-targeting antibiotics.

## Introduction

Cell-to-cell variation in isogenic bacterial populations is a common phenomenon that has been associated with cellular stress responses and anticipating potential environmental changes [1–3]. Heterogeneous survival to stressors such as antibiotics can occur through mechanisms including persistence, tolerance and heteroresistance, whereby heterogeneity may provide a survival advantage to a subpopulation of cells [4–6]. A potential strategy incurred due to heterogeneity is bet-hedging, where some cells prepare for a potential stressor in the future, enabling subpopulations of cells to resist once a stressor becomes present [7].

In contrast to permanent antibiotic resistance which is usually associated with genetic mutations, transient resistance is a temporary state which is lost once the stress is removed. Transient resistance is often associated with cell dormancy such as in persister cells, a subpopulation of cells which evades antibiotic killing, or in antibiotic tolerance, the ability of cells to survive high concentrations of antibiotic by slowing down an essential process such as growth [8–10]. RNA repair has been shown to promote transient antibiotic resistance via expression of the repair mechanism of the Rtc system, however, different to persistence, the transient resistance conferred by Rtc is characterised by growth rescue rather than dormancy [11, 12].

RNA, just like DNA, is susceptible to damage through mechanisms such as oxidative stress and radiation [13]. Damage may occur continuously through underlying cellular processes or infrequently through external stressors such as antibiotics [14]. Many RNA molecules have short-lived half-lives, such as mRNA in bacteria, and therefore a high turnover, meaning that damage can go largely unnoticed. However, some RNAs, such as ribosomal or transfer RNAs, are long-lived and damage to these molecules can have detrimental effects on cellular physiology [15]. RNA repair is therefore essential for repairing these molecules and maintaining the translational apparatus, hence ensuring cell growth and survival. The Rtc RNA repair system is highly conserved across all domains of life and is responsible for repairing damaged RNAs, which present as specific RNA ends, via the action of the repair proteins RtcA and RtcB [16, 17]. Repair systems like the Rtc system are inducible, and not constitutively active, meaning that single cells are able to sense and respond to damage accordingly. This mechanism allows cells expressing the Rtc system to rescue growth amid exposure to ribosome-targeting antibiotics, inferring a transient Rtc-induced resistance [11, 12].

Previous modelling revealed the heterogeneous nature of the RNA repair system through bistability and hysteresis at the population level, both in expression of repair proteins and in resulting damage control [12]. Bistability is a characteristic of deterministic systems which have two stable states, where the system can exist in either state depending on the initial conditions, therefore providing a description of distinct phenotypic states [18, 19]. Hysteresis is where the state of the system is dependent on its history [20]. However, both are properties of deterministic, population-level descriptions, and their interpretation at the single-cell level is not straightforward. It is not clear from population-level modelling how cells transition between different states or which state any given cell occupies at a given time [21, 22]. Bistability and hysteresis therefore reveal that heterogeneous Rtc-induced antibiotic resistance is possible, without explaining how it arises or per-sists at the level of single cells. Yet, understanding the drivers of state transitions is essential for understanding how cells respond to stress, and subsequently how they may survive in the presence of antibiotics.

Bacterial gene expression is a noisy, stochastic process which can be a source of substantial heterogeneity in mRNA and resulting protein levels between cells [23–26]. Variation in expression of genes related to antibiotic resistance has been shown to confer transient resistance in single cells which temporarily allows cells to survive antibiotic exposure [27]. A possible source of cell-to-cell variability in the Rtc system is the stochastic nature of Rtc gene expression itself [23]. The Rtc genes, *rtcA*, *rtcB*, both encoding repair proteins, and *rtcR*, encoding the regulator protein [16], are expressed at extremely low copy numbers meaning transcriptional activation is inherently a probabilistic event [23]. Stochastic effects matter a lot more in low copy number systems compared to high copy number systems, where continuum effects mean that stochasticity can often be more negligible [28]. Deterministic models are useful for understanding the behaviour of biological systems, but they do not account for stochasticity and therefore cannot capture variation across single-cell responses [21]. Stochastic models, on the other hand, are able to capture the inherent noise in individual cells and therefore capture the heterogeneity missed by deterministic models [29].

Despite previous work demonstrating the importance of RNA repair in response to antibiotics in *E. coli*, RNA repair has not yet been examined through the lens of stochastic heterogeneity. In this work, we develop a single-cell stochastic model of the Rtc system in *E. coli*, which we use to explore the effects of stochasticity on the responses of the Rtc system to RNA damage. We build upon our previous work by exploring what the deterministic characteristics such as bistability and hysteresis might look like in a stochastic regime. We observe bimodality in cellular translational capacity and identify a transient memory in response to RNA damage, consistent with hysteresis, suggesting that RNA repair promotes truly distinct phenotypes. To better understand the single-cell response to stress and the drivers of heterogeneity, we classify predicted cell trajectories to identify drivers of state switching between resistant and susceptible states. We find that stochastic expression of *rtcR* is the initial determinant driving the system into a resistant state, and dilution is the key driver of the system back into a susceptible state. RNA damage plays a role in both of these switching events, but it is not the main driver of either. Overall, these results demonstrate the importance of single-cell analysis to better understand heterogeneity and that variation in RNA repair in response to damage is driven by stochastic gene expression which may have implications for understanding how cells survive in the presence of translation-targeting antibiotics.

## Methods

### Modelling of RNA repair

In previous work, we developed a mechanistic model of the Rtc system [12]. Building on this, here we account for intrinsic stochasticity of gene expression and biochemical reactions. We couple the expression of the Rtc system and repair of a pool of translational RNAs to a deterministic cell cycle. We account for additional noise through division by binomial partitioning, enabling us to follow a cell lineage throughout a simulation. We implement the model in Julia (v1.11.5) using the Gillespie algorithm [30] with the `JumpProcesses.jl` (v9.21.2) package [31, 32]. The model is embedded within a package we created (`RtcModels.jl`) to ensure consistency in model versions with different users implementing the Rtc models. The package takes care of the model, associated parameters, initial conditions and includes code for simulation of the Rtc model. Simulations were performed using this package, which is available at https://github.com/hhindley/RtcModels.jl.

We implement a semi-stochastic cell division to allow for dilution in the model. The cell grows at a constant rate, λ, and once the volume doubles the contents are binomially partitioned between two daughter cells, one of which we keep simulating as in [33]. Division is performed by sampling a random number from a Beta distribution which dictates the proportion to be kept during binomial division. See SI for more details on the reactions and parameters used in the model. We perform simulations across a range of RNA damage rates (*k*_dam_) from 0.005 min^−1^ to 0.8 min^−1^ to observe the system under different levels of stress. For each damage rate we perform 2000 simulations for 1000 minutes each. We obtain two datasets in this way, one which is simulated using initial conditions obtained from a simulation run at low damage (0.005 min^−1^), and the other simulated with initial conditions obtained from a simulation run at high damage (0.8 min^−1^).

We include a method to initialise the model to account for transient phases when transitioning to different conditions. As we predominantly vary the damage rate, we firstly perform a preinitialisation simulation at zero damage to set pre-initial conditions. From this simulation, we randomly sample *n* = 2000 initial conditions for *n* initialisation simulations. To ensure the simulations are not still adjusting from the initial state and that there is variation in initial conditions for future simulations we then perform the initialisation simulations with these sampled initial conditions. We do these initialisation simulations at two different damage rates: low damage (0.05 min^−1^) and high damage (0.8 min^−1^), which gives us two datasets with different prior exposure to damage. We select initial conditions from our pool of initial simulations for the main simulations where we then sweep across damage rate (Figure S1).

### Hidden Markov models

We fit an HMM, using the Baum-Welch algorithm, to the healthy RNA concentrations for each damage rate individually on the data from simulations performed with previous exposure to low damage, as the simulations with prior exposure to high damage contained a broader range of data which could be explained with the extremes of the other dataset. We use the pomegranate package (v1.1.2) in Python to fit HMMs [34]. We fit three-state models to each damage rate where the three states represent susceptible, transition and resistant (Figure 1E). To obtain initial guesses for the emission distributions, we perform *k*-means clustering [35]. Once fitted, we constrain the transition state distribution to the means of the susceptible and resistant states, and the resistant and susceptible means to the means of each other to avoid unwanted misclassification in case data falls outside of the data range used to train the HMM. Emission distributions are described as a Gamma distribution for the susceptible state, a Uniform distribution for the transition state and a Normal distribution for the resistant state (Figure 1D) as this mixture of distributions gave the best log-likelihood across damage rates. We then use the Viterbi algorithm from `HiddenMarkovModels.jl` (v0.7.0) to determine cell states for each simulation.

**Figure 1:**
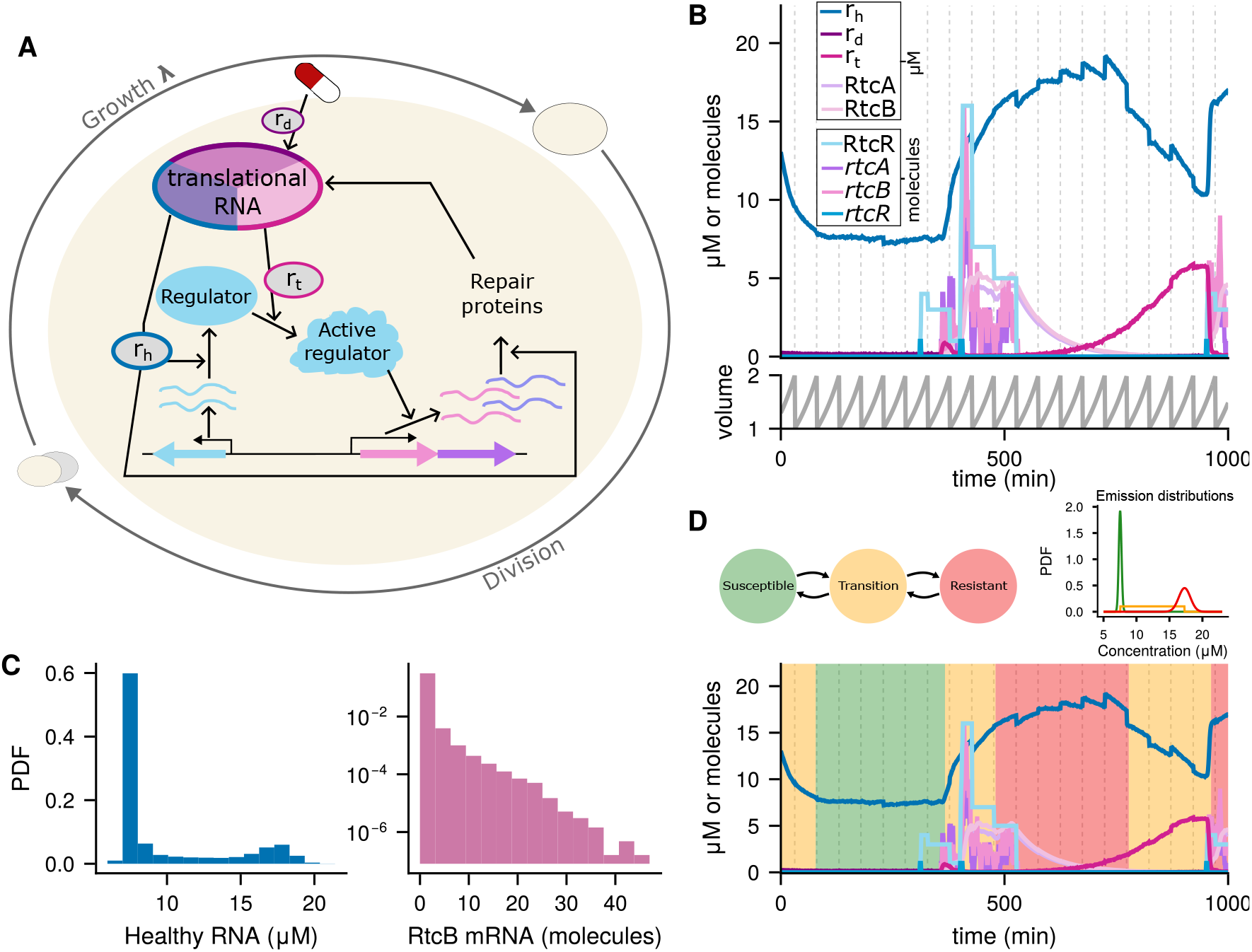
Modelling the Rtc system stochastically. (A) The stochastic Rtc model. Genes are transcribed to mRNAs which are translated to proteins by healthy translational RNAs (*r_h_*). The regulator (RtcR) is activated by tagged RNAs (*r_t_*) and active regulator activates transcription of the repair genes. The repair proteins (RtcA and RtcB), produced via translation by *r_h_*, repair damaged RNA *r_d_*, produced by indirect damage from antibiotics, to produce *r_h_* with *r_t_* as an intermediate species. The cell grows according to a growth rate λ and divides according to binomial partitioning once the volume has doubled (Methods). (B) A single simulation (cell) trajectory across time, at a damage rate of 0.02 min^−1^, shows all models species (top) and volume growth and division (bottom). Healthy RNAs increase in response to expression of the Rtc components and decrease in the absence of repair machinery. (C) Distributions of key model species: healthy RNA showing bimodality (left) and RtcB mRNA in log-y scale showing a zero-inflated distribution (right) for a damage rate of 0.02 min^−1^. (D) The structure of the three-state Hidden Markov Model model representing susceptible and resistant states with a transition state between them (top left). Emission distributions for the three states based on the fitted HMM for a damage rate of 0.02 min^−1^ (top right). The same simulation trajectory shown in C overlayed with HMM predicted states (bottom).

### Classification methods

To ensure results are consistent, we use two classification methods, logistic regression and random forest, to classify between cell state transitions. We only perform classifications on damage rates up to 0.4 min^−1^ to reduce the number of simulations needed to reach a reasonable sample size for classifications, furthermore, we observed a plateau or reduction in accuracy above 0.4 min^−1^. Logistic regression was performed as a ridge-regularized binomial classifier using the `GLMNet.jl` (v0.7.4) wrapper for the original GLMNet package in R [36]. We performed two classifications, the first classifying which state the system switched to after a susceptible to transition switch and the second classifying to which state the system switched after a resistant to transition switch. Each trajectory was split into 100-minute windows before both a susceptible to transition switch and a resistant to transition switch and labelled as either committing to the resistant state or to the susceptible state based on the state following the initial switch. Over each window we calculate a rolling Gaussian mean for each of the three mRNA species and each of the three protein species, giving us six features for each window. For classification of the next state given the susceptible to transition switch the classifier was trained on 6 samples per time point, with 2,224 positive and 4,369 negative examples and for classification of the next state given the resistant to transition switch there were 1,403 and 603 positive and negative samples respectively (damage rate 0.02 min^−1^. To ensure reasonable class sizes, we performed extra simulations on top of the 2000 simulations we performed for our base analysis (Table S3). Response labels were encoded as a categorical binary outcome.

Model evaluation used stratified 5-fold cross-validation with class balanced fold assignment to preserve the proportion of positive and negative examples across folds. Class weights were inversely proportional to class frequency to compensate for imbalance within each fold. Z-score standardisation parameters were estimated from the training data in each outer fold and then applied to the corresponding test set. Within each outer fold, GLMNet cross-validation was run on the training data to select the regularization strength that minimized mean loss, and the resulting model was evaluated on the held-out fold using predicted probabilities and ROC AUC. Coefficient extraction was performed by fitting the full standardized dataset with the same ridge logistic regression procedure, again selecting the best regularization parameter by cross-validation, and reporting the resulting coefficients for each species.

Random forest was performed using the `MLJ.jl` (v0.23.2) package [37] using a model with 200 trees, unlimited tree depth, a minimum leaf size of 2, and 3 randomly selected features considered at each split. For each damage rate, event-triggered windows were constructed in the same way as for logistic regression. As the random forest model in Julia does not implement class weights, the majority class was downsampled so that the two classes were balanced before model fitting. Model performance was estimated using stratified 5-fold cross-validation with the ROC AUC as the evaluation metric. Feature importance was obtained by fitting the random forest to the full dataset and extracting the built-in permutation-free feature importance for each species.

## Results

### Individual cells transition between distinct states leading to heterogeneity in cellular translational capacity

Building upon our previous deterministic mechanistic model [12], our stochastic model accounts for intrinsic noise in gene expression and repair actions, and includes cell division noise by implementing a simplified cell cycle. We assume cells grow and divide according to a constant rate λ, meaning division times are deterministic, and at division we keep track of one daughter cell by partitioning molecules binomially, therefore introducing noise to division events and tracking one cell lineage per individual simulation (see Methods). To account for the RNA repair actions of the Rtc system, we model a pool of three translational RNAs which represent RNA involved in translation. Healthy RNAs (*r_h_*) are the only species actively involved in translating mRNA to protein in the model, therefore they represent the translational capacity of the cell. Given translational activity drives cell growth [38], we consider translational capacity a proxy for cell susceptibility. In general, cells are considered to be susceptible to damage when there is a low concentration of healthy RNAs indicating a low translational capacity and resistant when there is a high concentration, indicating a high translational capacity. Damaged RNAs (*r_d_*) are those that have altered RNA termini caused by indirect antibiotic damage and are no longer functional for translation. Tagged RNAs (*r_t_*), which are also assumed non-functional, are produced by RtcA, which ‘heals’ the damaged RNA termini to produce RNA termini recognised by RtcR for its own activation. RtcB ‘seals’ tagged RNAs to completely repair the RNAs and reform fully functional translational RNAs.

Overall, the model contains nine model species: the three mRNAs expressed from the *rtc* operon, *rtcA*, *rtcB* and *rtcR*; the three proteins, repair proteins RtcA and RtcB and regulator protein RtcR; and the three translational RNA species, damaged (*r_d_*), tagged (*r_t_*) and healthy (*r_h_*) (Figure 1A, see SI for a detailed model description). We use the model to probe the effects of stochasticity on RNA damage and repair. We ask if single cells occupy distinct cellular states, to what extent they transition between states and what the primary drivers of these transitions are. Given we previously observed hysteresis in our deterministic model, here, we also characterise history-dependent responses in the stochastic setting.

We simulate individual cell lineages across different damage rates to observe resulting levels of stress on the system (Figure 1). Inspecting individual simulation trajectories, we observe a clear switching behaviour, where cells transition between high and low levels of healthy RNA, dependent on expression of the Rtc repair machinery (Figure 1B and C). We see the level of tagged RNA increase as healthy RNA decreases due to the rapid repair of damaged RNAs by RtcA producing tagged RNAs [39]. The repair action of RtcB is slower than that of RtcA [40], so there is an accumulation of tagged RNA before it gets converted by RtcB back to healthy RNA. We observe bursty expression of *rtc* genes to produce mRNA, and subsequently proteins, which precedes a switch to a high level of healthy RNA as the repair proteins convert damaged RNA back to healthy RNA (Figure 1B). Across many trajectories the switching between apparent states seen in the healthy RNAs manifests as bimodality (Figure 1C), where healthy RNAs either predominantly occupy a low state or a high state whilst there remains an underlying proportion between the two states denoting transitions between them. Bursty transcription of the Rtc repair genes yields a zero-inflated distribution (Figure 1C), similar to what we observed in single-cell experiments in our previous work and to the distribution expected from bursty gene expression [12, 41].

Next, for further analysis into state switching and to inspect history-dependent behaviour, we outline a method to predict states across our cell lineages. To classify translational states we use a three-state Hidden Markov Model (HMM) (Figure 1D). HMMs are probabilistic models used to predict a hidden, or unobserved, sequence of events given a sequence of observations over time. The hidden sequences are predicted from whole sequences rather than individual points, meaning a noisy observation is less influential than in other methods such as thresholding. We use healthy RNA concentration as the observation sequences for the HMM, as the trajectories span a range of concentrations and are always non-zero, therefore providing continuous sequences for state prediction. Each hidden state is associated with an emission distribution which describes the probability of observing a given RNA concentration conditional on the cell being in that state. Accordingly, our three-state model represents susceptible (S), transition (T) and resistant (R) cell states (Figure 1D). This is preferable over a two-state model (susceptible/resistant) as cells spend a substantial proportion of time transitioning between states, and a two-state model would result in overlap between the two states, or necessitate setting a threshold value between states. The use of an HMM in this instance, removes any potentially biased user input. States are instead classified according to the emission distributions predicted by the HMM (Figure 1D), using the Viterbi algorithm to determine the most likely state sequence. Our model predicts a low healthy RNA concentration to lead to the susceptible state, and a high concentration to represent the resistant state (Figure 1D). The presence of multiple states and switching between them produces an innate heterogeneity whereby genetically identical cells can be in different states at the same time leading to subpopulations of cells that are able to resist insult. Having observed state switching and bimodality, we ask whether there is any memory of previous state or condition in our system.

### Transient memory in response to RNA damage

In deterministic systems, memory of a previous state is known as hysteresis. In our previous work, we observed bistability, and associated hysteresis, in response to damage in our deterministic model of the Rtc system [12]. Now we introduce intrinsic stochasticity, we pose the same question: does memory exist in response to damage? Here, we observe the impact of initial conditions on system memory. To probe the dependence on initial conditions, we perform simulations where initial conditions are randomly sampled from prior simulations performed at a low damage rate, and those where initial conditions are randomly sampled from prior simulations performed at a high damage rate. Therefore, we have two sets of simulations initiated with previous low damage exposure and previous high damage exposure. To assess the differences between the two datasets, and to confirm the results are consistent across more than one metric, we compare distributions of healthy RNA using two distance measures: Jensen-Shannon (JS) distance and the Wasserstein distance. We choose to measure this distance between healthy RNA distributions and and not mRNAs, since the short half-life of mRNA means that any population-level information encoded in the mRNA distribution at *t*=0 is overwritten on the timescales relevant to our memory analysis. We initialise the system from the same set of initial conditions across different damage rates, one set for initialising with previous low damage exposure and one set for initialising with previous high damage exposure, allowing us to compare the temporal behaviour of the system under different values of our parameter of interest. As a result, at early time points the distances between distributions are identical across damage rates (Figure 2A).

**Figure 2:**
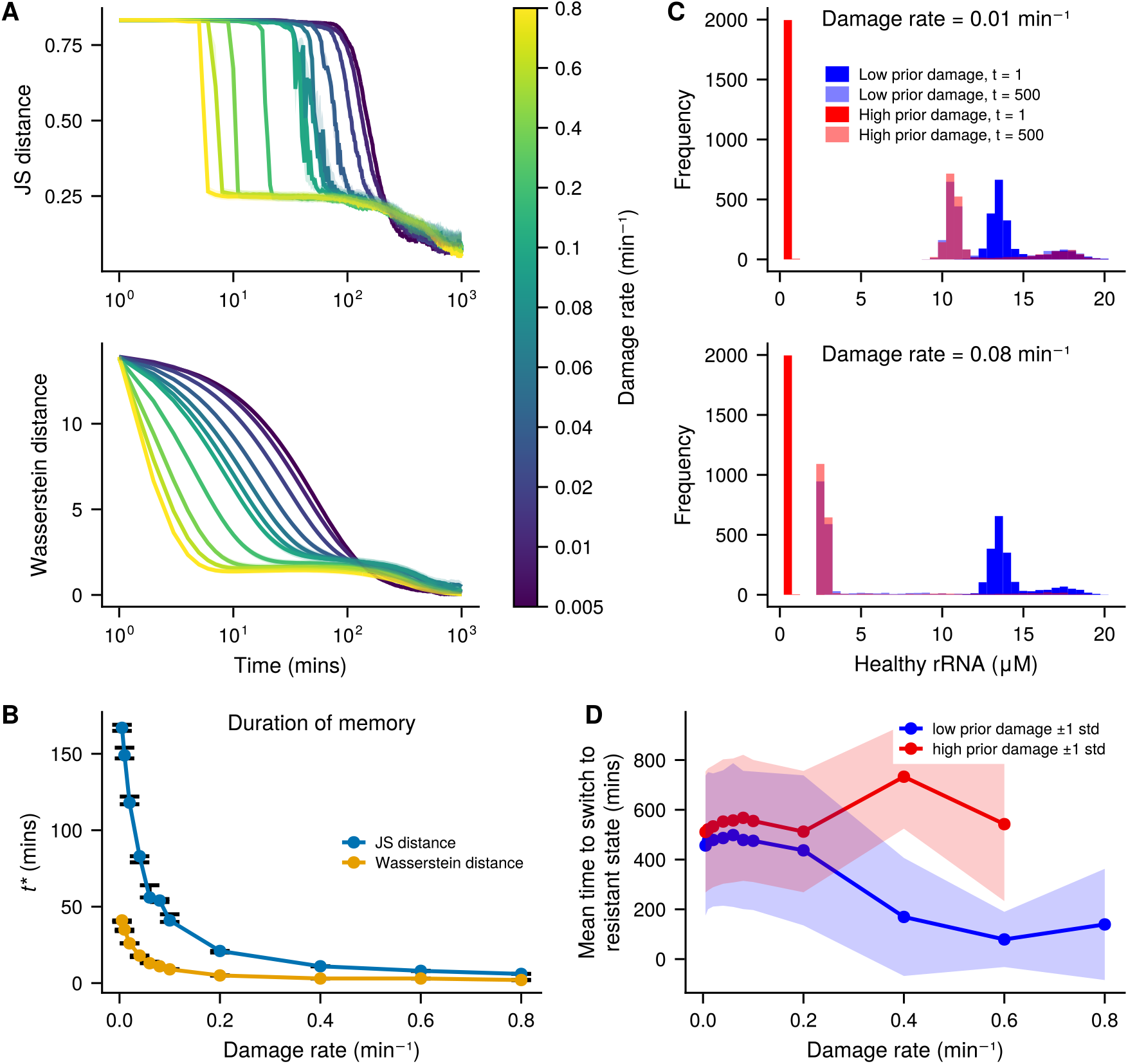
Transient memory in response to damage. (A) Distances between distributions initialised at low and high prior damage at every time point across damage rates. Top: Jensen-Shannon distance, bottom: Wasserstein distance. Shading around the lines represent the 95% confidence intervals from 1000 bootstrap samples. (B) The time taken for the distance between initialisation to reach half-maximal (*t\**) across damage rates. Error bars represent 95% confidence intervals across 1000 bootstrap samples. (C) Two sample damage rates (0.01min^−1^ top and 0.08min^−1^ bottom) showing the difference between distributions across simulations at time *t*=1 and *t*=500 for both initialisations. (D) The mean time taken to switch to the resistant state given the simulation started in the susceptible or transition states for both low prior damage initialisation (blue) and high prior damage initialisation (red). Shading around lines represents *±* 1 standard deviation.

Both metrics demonstrate eventual convergence of the two differently initialised distributions for all damage rates, suggesting that memory in this system is transient. The Wasserstein distance shows the distributions converging more gradually over time, and from the very first time point, compared to the JS distance, which predicts convergence to happen abruptly at a damage-dependent time point. Both metrics show that at low damage rates, the two initialisations diverge more markedly and for longer than at higher damage rates where they converge at earlier time points, demonstrating that initialisation has an effect on the temporal behaviour of the system (Figure 2A). Overall, the gradual convergence of the Wasserstein distance represents a more plausible scenario for distribution convergence, as the histograms are likely to change gradually across time rather than abruptly from one time point to the next.

We define *t\** to be the time at which the distance between the distributions is half-maximal to quantify the temporal relationship across damage rates, representing the duration of memory. We see that as the damage rate increases, the time taken for the distributions to converge decreases (Figure 2B). The two metrics differ in their *t\** values, with JS distance indicating a longer time for convergence across damage rates compared to the Wasserstein distance. This is reflected in Figure 2A where the Wasserstein distance decreases from the first time point whereas the JS distance remains constant at early time points, meaning the time for convergence is slower in the Wasserstein distance. This indicates that the initial conditions have a greater effect on the temporal behaviour at lower damage rates than at higher damage rates.

The difference in memory duration across damage rates can be explained by observing sample distributions at different time points and damage rates (Figure 2C). Simulations initialised with prior exposure to high damage have a lower initial starting concentration of healthy RNAs compared to those initialised from prior low damage exposure. Therefore, at early time points, the distributions between the two datasets are considerably different (Figure 2A and C). For the low and high prior damage initialisations to converge, the simulations from the high prior damage initialisation has to increase from this low initial concentration to be closer to the susceptible mode in the distributions from the low prior damage initialisation. As the damage rate increases, the concentration of healthy RNAs in the susceptible mode decreases, getting closer to the initial conditions of the high damage prior exposure dataset (Figure 2C). Therefore at higher damage rates, convergence to the susceptible concentration of healthy RNAs happens faster.

To further assess memory in our system, we observe the time taken to switch between states, specifically, the time taken to reach the resistant state, as the system is rarely initialised in this state. Simulations initialised with prior high damage exclusively begin in the susceptible state, as is expected due to high damage enforcing a low concentration of healthy RNAs. Simulations initialised with prior low damage mostly start in a transition state (Figure 2C). We take the simulations that started in either the susceptible or transition state and measure the time taken for the first switch to the resistant state, measured as the time point of the first occurrence of the resistant state. At low damage rates, the time taken to reach the resistant state is similar between low and high prior damage (Figure 2D). As the damage rate increases, the time to reach the resistant state in the low prior damage dataset decreases. However, the high prior damage dataset diverges away from low prior damage after a damage rate of 0.2 min^−1^. As damage increases, a prior exposure to high damage means cells are unable to readily switch to the resistant state. A damage rate of 0.8 min^−1^ prevents the high prior exposure dataset from entering the resistant state at all.

Overall, these results demonstrate a transient memory in the Rtc system in response to RNA damage, where the initial state has an effect on the temporal behaviour of the system and can dictate the rate at which the system enters a resistant state.

### Stochastic expression of mRNA drives the system into a resistant state

With switching displayed in the Rtc system in response to damage, we next aim to identify changes in model components which precede a switch and assess if this may determine state transitions. We are particularly interested in understanding the determinants of switching into the resistant state, as understanding this may suggest ways of preventing the resistant state occurring in the first place. After using the Viterbi algorithm to predict the states of the system, we identify switches from the susceptible to the transition state and align windows of time before these switches to observe the general behaviour preceding a switch (Figure 3A). We further separate these switching events into those that go on to switch to the resistant state (a full switch) and those that return to the susceptible state (an incomplete switch).

**Figure 3:**
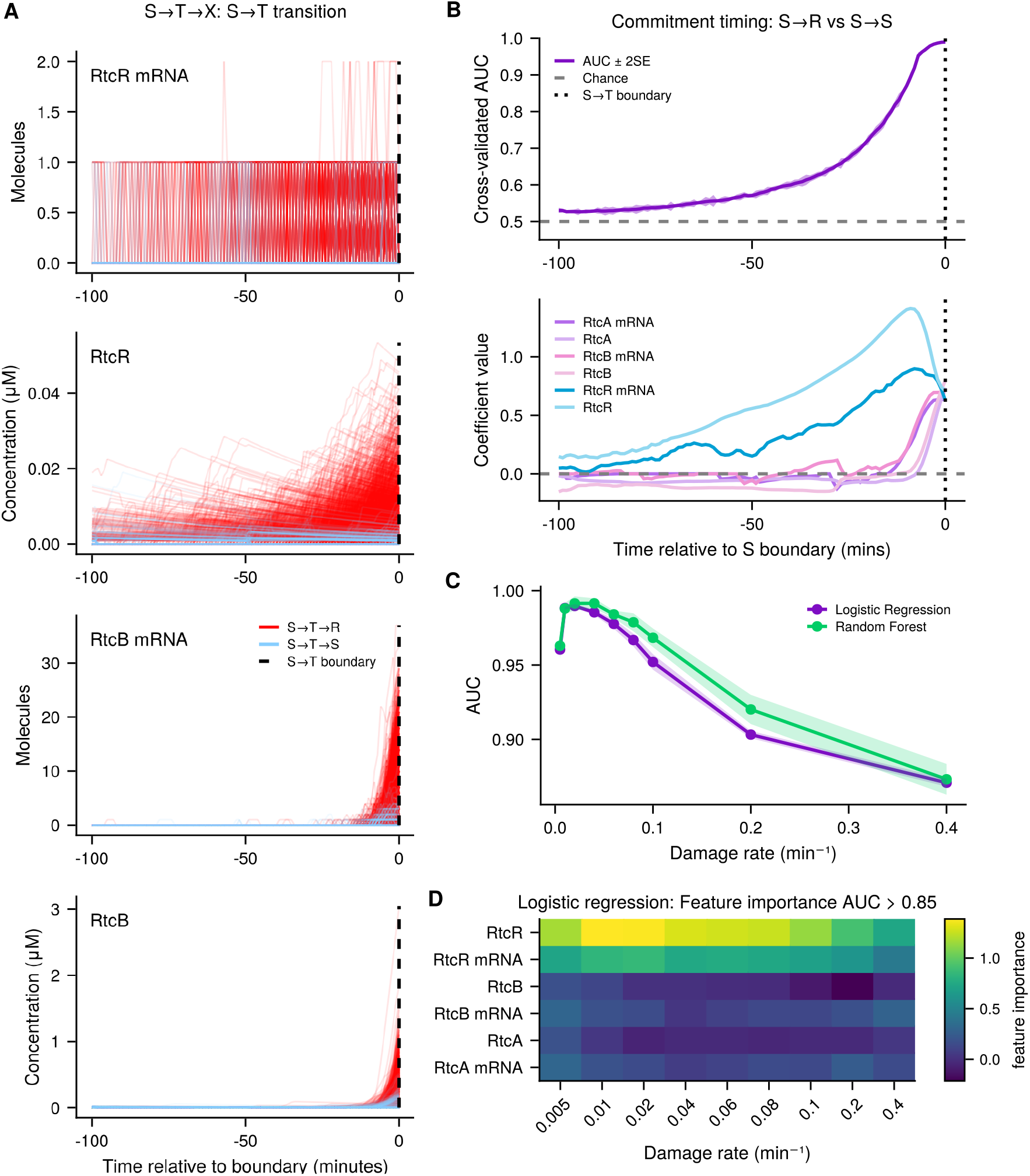
Identifying a driver of the resistant state. (A) RtcR and RtcB levels (mRNA and protein) during 100 minutes preceding a switch from the susceptible to transition state for RtcR and RtcB mRNA and protein. Red lines indicate full switches to the resistant state, blue lines indicate incomplete switches that return back to the susceptible state. (B) Logistic regression results for classifying between a full and incomplete switch. Top: AUC across time before a switch. Shading represents *±* 2 standard errors. Bottom: Coefficients for each species across time before a switch. (C) AUC across damage rates for both classification models. Shading represents *±* 2 standard errors. (D) Feature importance heat map across damage rates for logistic regression when AUC is higher than 0.85.

We observe an increase in all mRNA and protein species preceding a switch, starting with increased frequency of RtcR mRNA expression at around 40 minutes before a switch, followed shortly after by an increase in RtcR protein expression (Figure 3A). RtcB mRNA molecule counts and RtcB protein concentrations increase around 20 minutes before a full switch from susceptible to resistant (Figure 3A). We observe the same behaviour for RtcA mRNA and protein, as these are co-expressed alongside RtcB in the Rtc operon which we account for in our model. Rtc repair proteins are the final trigger for a full switch to occur, as they repair damaged RNAs to produce more healthy RNAs causing a switch to the resistant state. This dynamic analysis suggests that stochastic expression of RtcR mRNA is the initial driver of a switch to the resistant state. We exclude translational RNA trajectories from the full analysis since state classification is trained on the healthy RNA species, and the damaged and tagged RNA species are directly correlated with these, so their trajectories provide little independent information. We do not see any obvious activity for the incomplete switches, those that enter the transition state and then return to the susceptible state, suggesting that a lack of activity in these cases may be predictive of a switch back to the susceptible state.

To solidify the predictions made from our observations and to better identify important species for determining the fate of a switch we use two classification methods, logistic regression and random forests, to classify trajectories prior to switches by switching outcome, i.e. whether they go on to perform a full or incomplete switch. We give the classifier the mean of a 10-minute rolling Gaussian window for six species (mRNAs and proteins) across 100 minutes before the switch. For brevity, and because both methods show similar results, we only show the logistic regression results of one damage rate (0.02 min^−1^) for logistic regression (see Figure S2 for random forest results). Aligning with our qualitative analysis of trajectories (Figure 3A), the accuracy of the model, measured as area under curve (AUC), increases as we get closer to the switching time and remains highest in the last 20 minutes before the switch (Figure 3B). We further quantify feature importance by observing the logistic regression coefficients over time. As the accuracy rises the model predicts RtcR to be the most important feature for classifying between a full switch and an incomplete switch, with the corresponding mRNA also being important, suggesting that the model can predict a switch to the resistant state without the expression of repair proteins (Figure 3B). In the last 20 minutes before a switch, when the accuracy is at its highest, the coefficients of the repair proteins steeply increase, indicating that the model can confidently make the correct prediction of a switch once the expression of repair proteins has occurred (Figure 3B). Given the protein expression of RtcR must be preceded by *rtcR* gene expression and that *rtcR* expression is a constitutive, purely stochastic event that does not rely on any other model species or dynamic variables, we can assume that resistance occurs, in the first place, due to stochastic transcription events of *rtcR*.

Interestingly, with both classification methods, the accuracy of classification decreases as damage rate increases (Figure 3C). At high damage rates the system requires a big burst of RtcBA expression to reach the resistant state, however, if the burst is not strong enough, specifically if there is not enough RtcB to convert tagged RNAs back to healthy RNAs, then the system will return to the susceptible state as damage continues to occur. This means that at high damage rates there is expression of mRNAs and proteins before a switch from the susceptible to transition state even in the incomplete switch case, however, switching outcomes are more uncertain, which makes it harder to classify between the two cases and reduces the accuracy of the model (Figure S3).

We observe that RtcR being most predictive of a switch holds across damage rates (Figure 3D), where logistic regression predicts RtcR and the mRNA to be the most important features for classifying with an AUC above 0.8. Random forest classification predicts RtcR to be the most important feature across damage rates, with less of an emphasis on the importance of the RtcR mRNA (Figure S4). The importance of RtcR in determining a switch to the resistant state aligns with previous work showing that inhibition of RtcR might inhibit the Rtc system and reduce cellular translational capacity [12].

Although entering the resistant state is driven by stochastic expression of RtcR, which does not depend on damage, the damage rate will modulate how often these bursty expression events translate to a switch into the resistant state: at higher damage rates, the fraction of active RtcR, caused by the interaction of RtcR protein with tagged RNA molecules, will be higher due to a high damage producing a higher number of damaged RNAs which are subsequently converted into tagged RNAs by RtcA. According to our previous model [12], we include a baseline level of leaky activation of RtcR in the model to account for the fact that RtcR activation requires the binding of a ligand in addition to the presence of both RtcA and RtcB [42]. Furthermore, this means that in the absence of RtcB or RtcA, any RtcR present will be activated at a low level. Leaky RtcR activation allows for some expression of RtcBA for initiation of a positive feedback loop in the model (Figure 1B) that ultimately leads to a switch to the resistant state. This means that at higher damage rates, the system is more likely to switch to the resistant state given a burst of RtcR expression, as there will be more active RtcR available to initiate positive feedback.

### Dilution drives the system into a susceptible state

As well as understanding how cells enter the resistant state, we would also like to understand how they leave the resistant state and transition to the susceptible state. Using the same approach as above, we align trajectories and consider dynamics during 100 minutes preceding a resistant to transition switch and again distinguish between full switches, those that transition on to the susceptible state, and incomplete switches, those that return to the resistant state. Unlike before, time windows here are devoid of a significant signal predictive of a switch to the susceptible state. For a damage rate of 0.02 min^−1^, full switches show a slight decline in the proteins as the window approaches the switch (Figure 4A) and incomplete switches show a mixture of species decline and mRNA and protein expression. Other damage rates display similar behaviour, but we see a more substantial difference in full and incomplete switches at lower damage compared to higher damage rates (Figure 4A vs S3B). Once repair protein levels can no longer sustain repair against the damage rate present, the system begins accumulating damage and exits the resistant state. This exit happens faster at higher damage rates, so full and incomplete switches become harder to distinguish in the 100-minute window preceding a switch. Both full and incomplete switches display comparable protein and mRNA levels remaining from the preceding resistant state, as well as from new expression leading to an incomplete switch (S3B). At lower damage, the system takes longer to leave the resistant state once repair protein levels drop below a sustainable threshold, so the pre-switch window shows correspondingly less expression activity for those cases switching to the susceptible state. By contrast, incomplete switches display some expression activity from new expression events, making these more easily distinguishable from full switches at low damage.

**Figure 4:**
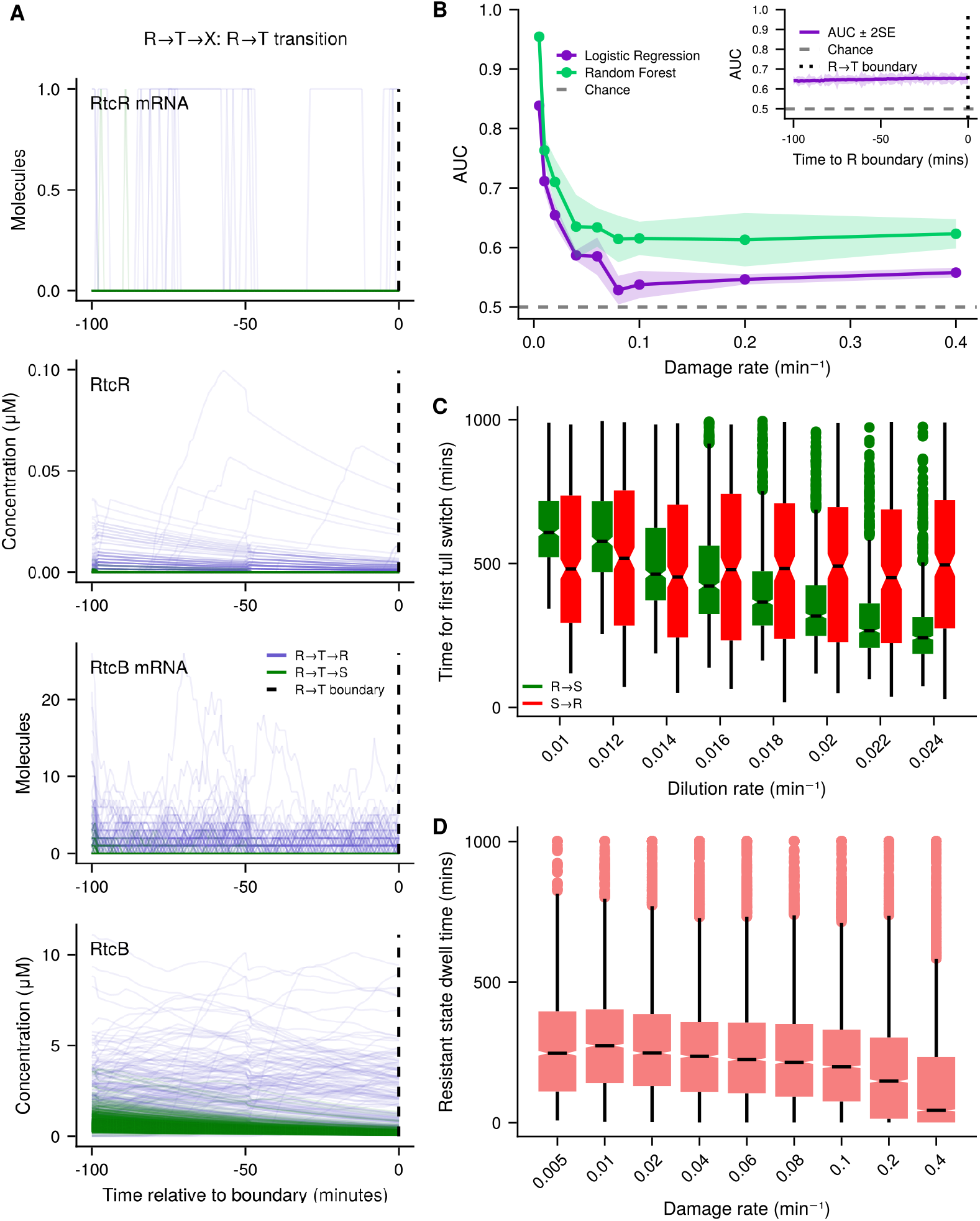
Identifying a driver of switching out of the resistant state. (A) Time windows preceding a resistant to transition switch for RtcR and RtcB mRNA and protein. Green lines indicate full switches to the susceptible state, blue lines indicate incomplete switches that return to the resistant state. (B) AUC across damage rates for both classification models. Shading represents *±* 2 standard errors. Inset: damage rate 0.02 min^−1^ logistic regression accuracy over time. Shading represents *±* 2 standard errors. (C) Box plots of time taken to make the first full switch from simulation initialised in the resistant state (blue) and the susceptible state (green) across different dilution rates. (D) Box plots of dwell times in the resistant state across damage rates. Box plots highlight median, interquartile range (IQR), with whiskers spanning the third and first quartiles *±* 1.5 x IQR.

As before, we then classify between full and incomplete switches. Accuracy of the classification model is highly dependent on the damage rate as AUC decreases sharply until it plateaus from around 0.1 min^−1^ where it is approximately 0.54 *±* 0.01 and 0.62 *±* 0.01 for the logistic regression and the random forest, respectively (Figure 4B). Differences in accuracy across damage rates may be due to our observations stated above, where low damage rates display a stronger contrast between full and incomplete switches, thus enabling easier classification compared to high damage rates. We also note that the AUC does not increase for time points approaching the switching time as it did for the susceptible to transition switch, but instead remains constant (Figure 4B inset). At low damage rates, classification is driven by expression of repair proteins being indicative of incomplete switches, those that return back to the resistant state (Figure S5), consistent with our previous analysis where we observed an increase in repair proteins just before a switch out of the susceptible state towards the resistant state. This also tells us that in order to transition all the way to the susceptible state, the system has to drop below a threshold of repair proteins. However, this classification result is not informative about which species drive the system to the susceptible state.

These results indicate there is no species truly indicative of a switch out of the resistant state, as logistic regression did not capture a positive coefficient for any feature when the model performed at high accuracy. This, along with the fact that trajectories seem to be decreasing across the windows preceding a switch, suggests switching out of the resistant state could be driven by dilution or damage.

To test the dilution hypothesis, we run initialisation simulations across different dilution rates, we fit an HMM to each dilution rate and predict the states given the healthy RNA concentration. We take a random sample of initial conditions from only resistant states and use these for new simulations across different dilution rates. We then observe the time taken to switch to the susceptible state from the resistant state. As dilution rate increases, the time taken to switch to the susceptible state decreases (Figure 4C). This suggests that dilution impacts switching back to the susceptible state, and indeed affects how long the system stays in the resistant state. We perform the same analysis measuring the time it takes to transition initially from the susceptible state to the resistant state, and corresponding to our above analysis we find the dilution rate does not influence the time taken to switch to the resistant state from the susceptible state (Figure 4C). We quantify the differences in distributions of first switch time between dilution rates with *p*-value testing, using pairwise Mann-Whitney-U tests with Bonferroni correction, and confirm that the differences between most adjacent dilution rates are significant for exiting the resistant state and that most are not significant when exiting the susceptible state (Figure S6).

Given that a switch out of the resistant state occurs when the concentration of healthy RNAs decreases below a certain threshold, the damage rate may also modulate this transition. We observe a decrease in resistant state dwell times as damage rate increases (Figure 4D), suggesting the damage rate affects the switch out of the resistant state. The system will not leave the resistant state if there are sufficient levels of repair proteins available to counteract damage. Therefore, to leave the resistant state, the repair proteins have to be at a low enough concentration for healthy RNAs to be decreasing due to damage. Once this happens, the concentration of healthy RNAs will decrease due to both damage and dilution, eventually reaching a low enough concentration to enter the susceptible state, given there are no new repair proteins formed. Therefore, the first stage of exiting the resistant state is dilution of repair proteins. Following this, a combination of dilution and damage drives the system out of the resistant state. Overall, these results suggest that dilution is the first driver of a switch back to the susceptible state, but like a susceptible to resistant switch, this switch is also modulated by the damage rate.

## Discussion

In this work, we present a stochastic model of the Rtc RNA repair system. The model considers the expression of three *rtc* genes, whereby corresponding RNA repair proteins modulate a pool of translational RNA species. We consider the damage rate to be a proxy for stress on cells triggered by translation-targeting antibiotics and we regard RNA damage to occur via the indirect action of antibiotics producing oxidative stress [43]. Damage has a direct effect on the concentration of healthy RNAs, the species which determines cell state, and on the expression of repair proteins. We systematically simulate the model using the Gillespie algorithm to analyse how stochastic effects dynamically drive cell fates between resistant and susceptible states. We demonstrated that the damage rate modulates switching between these states, therefore, suggesting the concentration of antibiotics could influence whether subpopulations of Rtc-induced resistant cells form or not, which could substantially impact the effectiveness of antibiotic treatments.

Our previous work on a deterministic model of the Rtc system revealed bistability. In a deterministic system, bistability allows for two coexisting stable states, and hysteresis directly follows, meaning that once a trajectory settles into one steady state or basin of attraction, it remains there, retaining permanent memory of its initial condition. A stochastic system, however, is continually subject to fluctuations. Therefore, in our stochastic setting we observe a transient memory, the rate at which simulations from low and high damage initialisations converge towards the stationary distribution is history-dependent. Essentially, the transient regime is what determines a cell’s effective phenotype. We therefore suggest that bimodality, the presence of switching behaviour, and a transient memory constitute the appropriate stochastic counterparts to deterministic bistability and hysteresis, and that characterising these signatures, rather than searching for permanent hysteresis, is the correct framework for identifying bistable-like behaviour in stochastic systems.

We analyse heterogeneity in the Rtc system quantified according to the translational capacity of cells, measured as the concentration of healthy RNAs available for translation. This heterogeneity displays as bimodality in healthy RNA concentration, where cells mostly occupy one of two states, a high concentration of healthy RNAs, corresponding to cells with a high translational capacity and therefore a resistant phenotype, or a low concentration of healthy RNAs, corresponding to cells with a low translational capacity and therefore a susceptible phenotype. Cells residing in between these two states are in a transition state and can transition to either of the susceptible or resistant state. In our system, the transient nature of this memory is modulated by the damage rate. Previous states are more important in determining the future behaviour of the system at low damage rates as different initialisations converge slower compared to high damage rates. This may have implications for treatment strategies, where initially exposing cells to a high damage, meaning a higher antibiotic concentration, may mean that individual cells are less likely to launch repair and thus subsequent treatments are more successful.

We show that stochastic expression of *rtcR* is the first essential component of switching to the resistant state, and that dilution and damage are the prominent drivers of returning back to susceptibility. This demonstrates the potential significant effects of stochastic gene expression, as, given the right conditions, it can push cells into a resistant state. A low dilution rate, or growth rate, means that the system can persist in the resistant state for longer, so higher growth rates are potentially beneficial for treatment strategies. The growth rate of cells is known to be affected by the presence of antibiotics, where cells may enter a dormant state and slow their growth rate to survive such as in persister cells. This may mean that cells are able to persist in the resistant state for longer, and therefore survive antibiotic treatment.

A key limitation of our model is that the growth rate is not dynamic and linked to the concentration of healthy RNAs. We made this modelling choice for simplicity and to isolate the Rtc system from other cellular processes. However, in reality, the growth rate has been shown to be directly proportional to the concentration of cellular ribosomes, raising interesting questions for future investigation. Secondly, we do not model the translation-inhibiting effects of antibiotics, under which growth rate and ribosomes are negatively correlated. Instead, we focus on the repair to investigate how this can promote heterogeneity, and note that more work will be required to fully incorporate these antibiotic effects. Finally, whilst our model parameters mostly come from reputable literature sources, some parameters are afflicted with high uncertainty. We previously performed a sensitivity analysis to address this in our deterministic setting, whereas the focus here was to characterise the nature of heterogeneity and identify drivers of cell state transitions [12].

Overall, we have demonstrated here the importance of RNA repair in response to antibiotics, and how stochasticity in gene expression promotes heterogeneity in the cellular translational capacity modulated by the Rtc system. We have identified key drivers of switching behaviour, of which results align with previous findings that RtcR inhibition may be effective at preventing the onset of the resistant state [12]. We outline the importance of growth rate in determining how long cells persist in the resistant state, and therefore a potential strategy for surviving antibiotic treatment. In view of rising levels of antibiotic resistance, our predictions provide testable hypotheses to unravel a deeper understanding of bacterial responses to a widely-used class of antibiotics and thus pinpoint avenues to potentiating their use.

